# A Comprehensive *In Silico* Study for the Identification of Therapeutic Target Against Peripheral Neuropathic Pain in humans

**DOI:** 10.1101/2021.11.03.467110

**Authors:** Sagar Bhowmik, Sheikh Mohiuddin Samrat, Debneela Paul

## Abstract

**Background:** VGF (non-acronymic) is a neuropeptide precursor or neuro-protein or neurosecretory protein which plays vital roles in the regulation of gastric contractility, mood regulation, and peripheral neuropathic pain and possibly, cancer.

**Objective:** VGF may be a potential target as it has a unique contribution to the development of neuropathic pain which is a target for Oxymatrine (OMTR).

**Method:** Based on this, we have endeavored to discover VGF inhibitors from the ChEMBL database of Oxymatrine (OMTR) analogues by employing homology modelling, molecular docking and pharmacophore analysis.

**Result:** Our *in silico* investigation reveals that 13-Methoxymatrine has desired characteristics for becoming a future formulation.

**Conclusion:** To confirm the efficacy of this compound, essential animal and clinical trials are needed to be performed. We believe that our present study will help to find an efficient and effective therapy for treating neuropathic pain in human which is modulated by VGF.

## 1. INTRODUCTION

VGF (nonacronymic) is a neurotrophin-inducible neuropeptide precursor that has potent antidepressant-like actions and regulates energy metabolism [1–5], two actions that might be related. VGF, expressed abundantly in neurons and some neuroendocrine cells, is cleaved by proteases into smaller peptides like TLQP-21, TLQP-62 that are subsequently secreted. Due to the presence of paired basic amino acid residues (R — Arginine, and K– Lysine), the VGF sequence undergoes endoproteolytic cleavage to produce several smaller peptides, released upon stimulation via the regulated secretory pathway both in vitro and in vivo. There are data suggesting that the VGF-derived peptides are the biologically active, stored in dense core vesicles and secreted in order to play a role in inter cellular communication, and responsible for the diverse range of biological functions associated with VGF like energy balance, re-productive behavior, pain modulation, mood order, etc. [1, 6]. Heat shock cognate protein A8 (HSPA8) has been found as a receptor of VGF derived bioactive peptide TLQP-21 in SH-SY5Y cells.

So, it was of much interest to observe whether Oxymatrine (OMTR), an inhibitor of HSPA8, inhibits the binding of TLQP-21 to the surface of live SH-SY5Y cells. The results confirmed, as expected, that OMTR reduces the binding of TLQP-21 to the surface of live SH-SY5Y cells, with a strong conclusion that the binding of TLQP-21 to the surface of SH-SY5Y cell model was through HSPA8 [7, 8]. HSPA8 (71 kDa), a constitutively expressed protein, is a fascinating member of the HSP70 family. HSPA8 expressing on the cell surface performed as a cellular receptor [7, 8]. Over 70 candidate compounds were screened for HSPA8 inhibitor. Among the compounds examined, Oxymatrine (OMTR, matrine oxide, matrine N-oxide, matrine 1-oxide: one of many quinolizidine alkaloid compounds), molecular weight (MW) 264.31, an alkaloid extracted from *Sophera flavescens* [9] (popularly known as KuShen plant), showed significant activity to down regulate the expression of HSPA8 in HepG2 liver cells without showing any toxicity to the cells. Additionally, Western blotting confirmed the reducing effect at the protein level, showing reduction of HSPA8 protein in the OMTR treated cells [10]. Use of active site-targeting inhibitor is an effective approach leading to pharmacological inventions. The inhibition efficacy of OMTR potentiates its application in drug targeting. On the other hand, VGF contributes to the modulation of pain through activation of primary microglia and potentially involve interactions with components of the complement system. The findings highlight the importance of VGF in the research of conception and could lead to new perspectives and targets for pain therapeutics. Considering all these, the study here was undertaken to ascertain VGF inhibitors from the ChEMBL database of Oxymatrine (OMTR) analogues *in silico*.

## 2. METHODOLOGY

### 2.1. Homology Modeling, Model validation and Prediction of Active Site

The protein primary sequence of VGF (vascular growth factor) was retrieved from NCBI database [11] and NCBI-BLAST (Basic Local Alignment Search Tool) [12] was used for the identification of suitable templates for homology modeling of VGF. The homology model was built by EasyModeller ver 4.0 [13], SWISS-MODEL [14–16] and CPH MODEL [17]. The generated models were validated by VERIFY3D server [18] which uses three-dimensional profiles for assessing protein models and SWISS-MODEL workspace of ExPASY server [19] for PROCHECK [20] and QMEAN6 (Qualitative Model Energy Analysis) [21] value. PROCHECK assesses the stereochemical quality of a protein structure and QMEAN stands for Qualitative Model Energy Analysis, is a composite scoring function describing the major geometrical aspects of protein structures. After analyzing the data from server, the best suitable model is selected. The modeled structure and templates were superimposed by UCSF Chimera ver 1.12 [22]. The active site residues which will play an important role in protein-ligand interaction were identified by the COACH server [23].

### 2.2. Virtual screening for the retrieval of potentially active compounds

ChEMBL server [24] which is now a well-established resource in the fields of drug discovery and medicinal chemistry research. The ChEMBL database curates and stores standardized bioactivity, molecule, target and drug data containing 2,101,843 compounds. This server was employed for finding structural analogous of Oxymatrine using 70% similarity threshold search and 22 compounds were found. These compounds were submitted to SwissADME server [25] and screened based on Ghose violations [6, 26–29]. After the primary screening, all of the compounds were submitted to PASS (prediction of activity spectra for substances) prediction server [30] which predicts the possibility of active (Pa) and inactive (Pi) value for identifying potent compounds which have antimycobacterial activity. Based on Pa and Pi value from PASS server and toxicity profile (mutagenic, tumorigenic, reproductive, irritant effect) in accordance with a good drug-likeness score by DataWarrior software [31], 5 leads were selected for further study.

### 2.3. Molecular Docking

The selected ligands and protein model were optimized by UCSF Chimera ver 1.12 [22] before docking was performed. Docking was performed by AutoDock Vina ver. 1.1.2 [32] which predicts the interaction between ligands and bio macromolecular targets. Before docking, polar hydrogens were added to the Protein model of VGF by Autodock tools and initial parameters were assigned for the protein molecule. For the compounds, the torsions were fixed by Autodock tools and both the protein and compounds were saved as PDBQT file format. The grid (grid size -98*92*78) was fixed around the active site which was obtained from COACH server. On 6 different poses, the binding energy of the protein and ligands were obtained and the best binding energy was chosen for further analysis. LIGPLOT [33] was employed for presenting the two-dimensional interaction between protein and ligands. Pymol [34, 35] was employed for presenting the three-dimensional interaction between protein and ligands.

### 2.4. Molecular properties and Oral toxicity prediction

The pharmacokinetics, ADME and toxicity properties of the lead compound were estimated using SwissADME [25] and PreADMET [36].

## 3. RESULT and DISCUSSION

CADD or computer-aided drug discovery requires to perform three main tasks, namely, prediction of the three-dimensional structure of the protein, prediction of the possible interaction between protein and ligands by docking and finally, testing biological and pharmacophore properties of the lead which was targeted as potential drug [37]. The following is performed in this study to carry out the research.

### 3.1. Homology Modeling, Model Validation and Prediction of Active Site

The absence of three-dimensional (3D) structure of VGF (vascular growth factor) in the public database, instigated us to build a 3D structure using homology modelling method by SWISS-MODEL, CPH Model and EasyModeller. From analyzing the data from PROCHECK, VERIFY3D and QMEAN6; model generated from EasyModeller is proved to the best suitable for our purpose. EasyModeller uses two templates (3j83.1A) and (2I1J. A) which have 34.62% and 26.47% sequence identity respectively. Subsequently, VERIFY3D server showed that the modelled structure had 48.48% residues had an averaged 3D-1D score >=0.2. PROCHECK showed that 87% and 9.2% residues in the most favored and additional allowed region respectively according to Psi (degrees) and Phi (degrees) and 1.6% residues in the disallowed region in the Ramachandran Plot Figure 4. QMEAN6 showed that z-score is -10.12 Figure 5. COACH server predicted 10, 200, 204, 208, 211, 242, 245, 246, 293, 296, 314,315, 318, 319, 321, 335, 340, 345, 353, 356, 359, 363, 365, 366, 367, 368, 376, 405 and 408 residues as potential active site residue. For designing a drug, homology modeling is a reliable as well as quick method for obtaining a protein model which is currently unavailable in the present databases [38–40]. It is opined that if any proteins don’t have structural information then the model of that protein can be built by homologous proteins or templates which have at least 30% sequence identity [41]. In this study, the templates have more than 30% sequence identity Table 1 which is highly reliable for constructing a model based on the template.

**Table 1.**
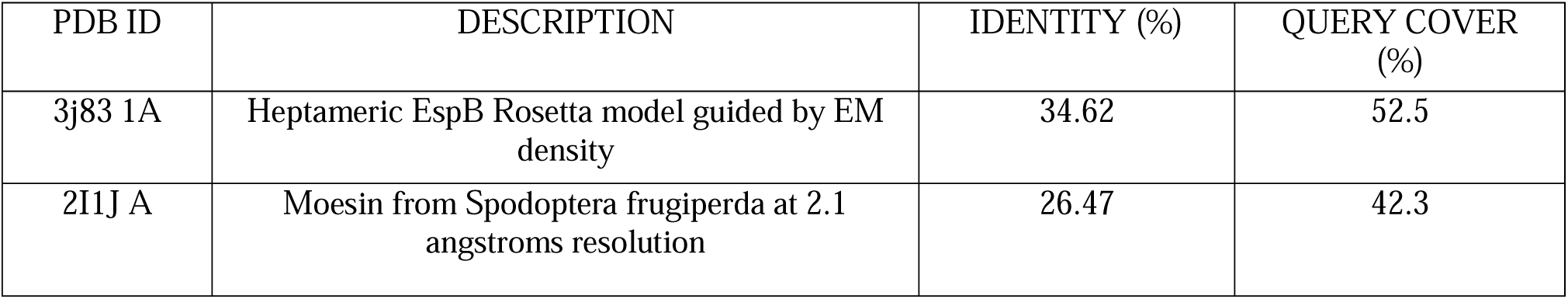
Templates used for homology modeling of VGF.

**Table 2.**
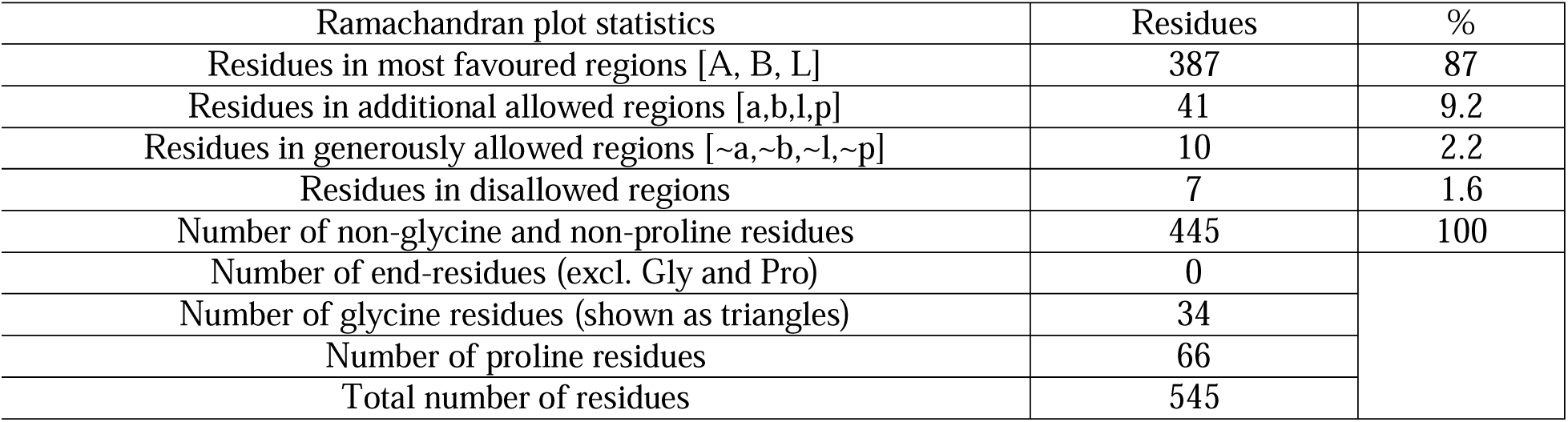
Ramachandran plot statistics for generated protein model.

**Table 3.**
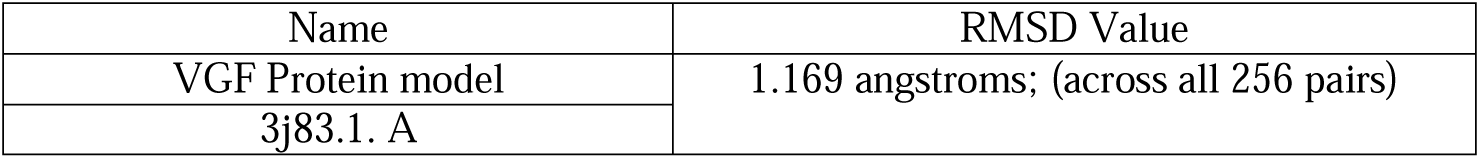
Root mean square deviation (RMSD) between VGF model and template (3j83.1. A).

**Table 4.**
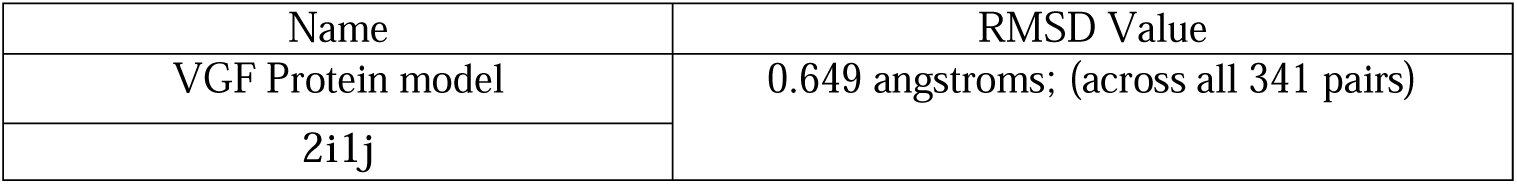
Root mean square deviation (RMSD) between VGF model and template (2i1j).

| Name | RMSD Value |
| --- | --- |
| VGF Protein model | 0.649 angstroms; (across all 341 pairs) |
| 2i1j |  |

From Ramachandran Plot and VERIFY3D, it is transparent that the built model was valid. Additionally, in QMEAN6 validated the modeled structure based on six structural descriptors. The modeled structure and the superimposed structure of generated model and template showed that the model was correct. The modeled structure and the superimposed structure of generated model and template are shown in Figures 1, 2, 3, 6, 7 and 8.

**Fig. 1.**
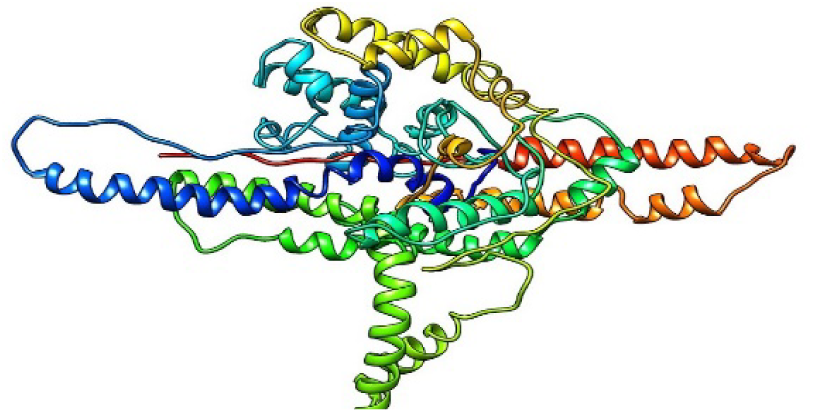
The modeled protein in ribbon style.

**Fig. 2.**
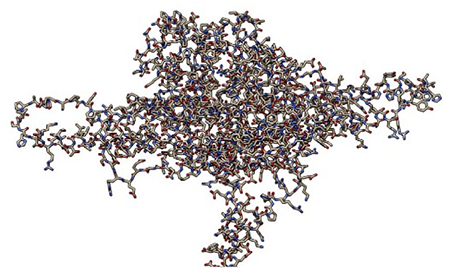
The modeled protein in wireframe style.

**Fig. 3.**
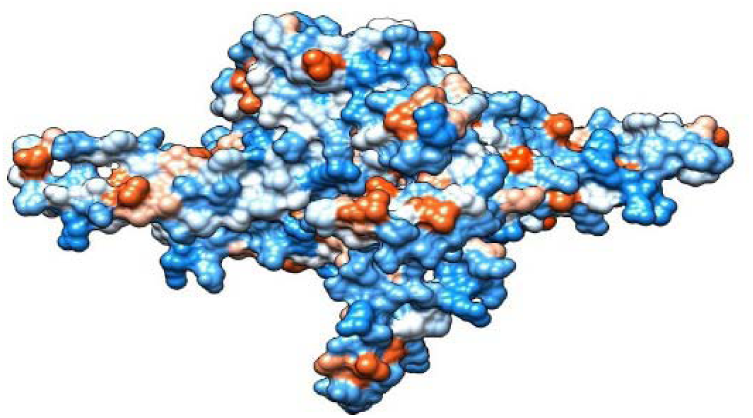
The modeled protein in hydrophobicity surface style.

**Fig. 4.**
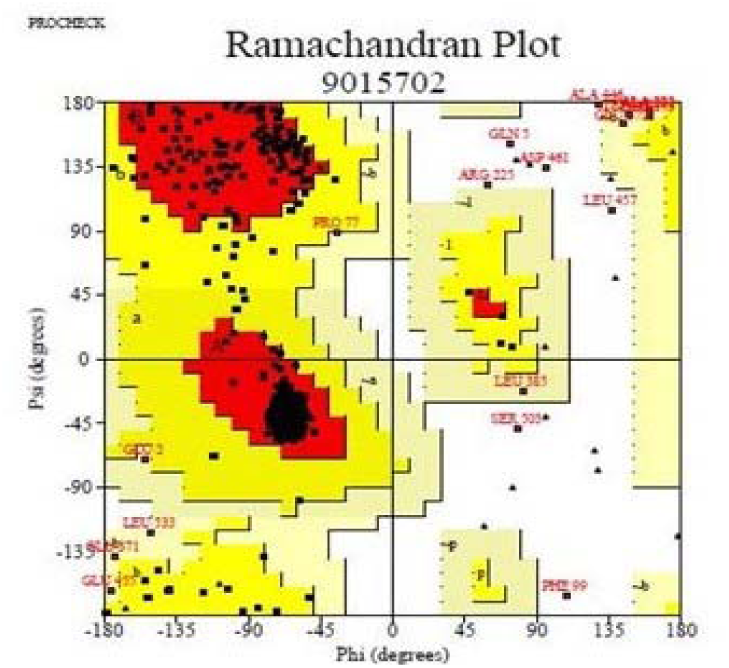
Ramachandran plot of modeled structure validated by PROCHECK server.

**Fig. 5.**
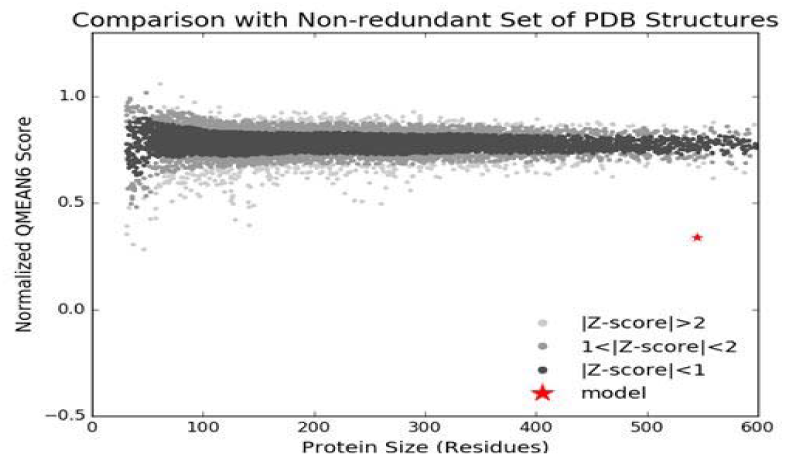
Validation of the modeled structure by QMEAN6.

**Fig. 6.**
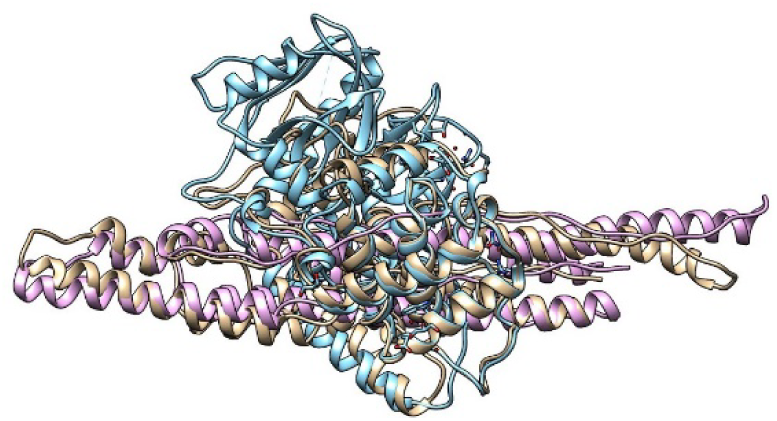
Superimposed structure of the modeled protein and templates.

**Fig. 7.**
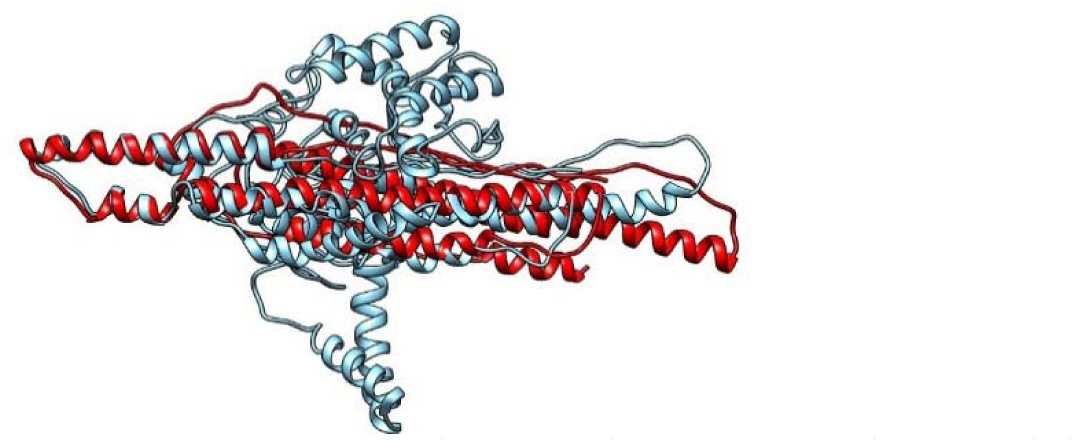
Superimposed structure of the modeled protein (blue color) and template (3j83.1. A) (red color).

**Fig. 8.**
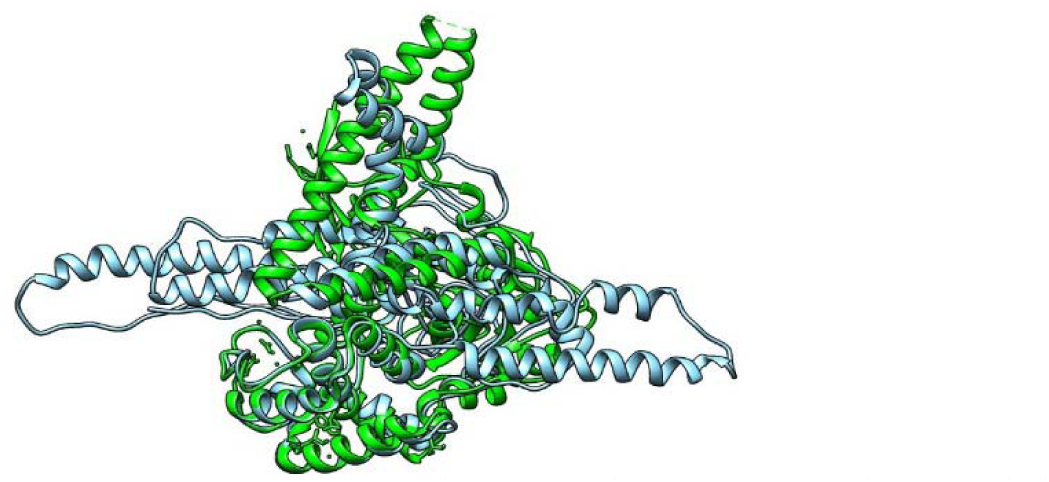
Superimposed structure of the modeled protein (blue color) and template(2i1j.pdb) (green color).

**Fig. 9.**
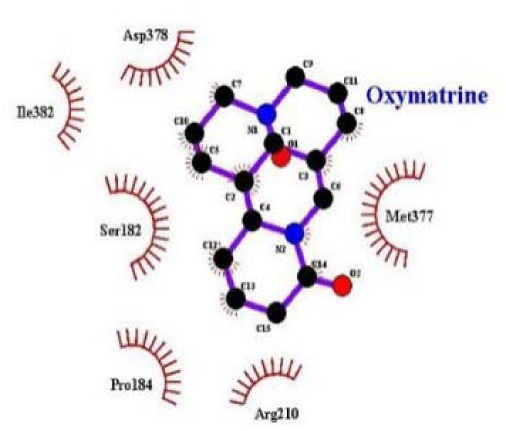
2D interaction between protein and Oxymatrine.

### 3.2. Virtual screening for the retrieval of potentially active compounds

Based on similarity search in ChEMBL, 22 compounds were found among which 7 compounds followed Ghose rule, Lipinski rule, Veber rule, Egan rule and Muegge rule and selected for future work. Among these 7 compounds, 6 compounds which had a good antimycobacterial activity were selected based on higher Pa value compared to lower Pi value from PASS prediction server. Compounds which didn’t show any mutagenic, tumorigenic, reproductive, irritant effect and had a good drug-likeness score were finalized as leads and by this, 5 leads were found.

ChEMBL is an Open Data database which contains more than one million compounds especially for the drug discovery and chemical biology research [24]. From this database, 22 compounds of Oxymatrine were primarily screened which followed five rules which are used for screening leads currently. Oxymatrine was selected as the mother ligand because of this ligand functions as the inhibitory agent against VGF [7] and this neuropeptide VGF (a non-acronymic name) is responsible in the regulation of neuropathic pain in human [42]. Ghose’s rule [26, 43] explained that a compound should have calculated log P between *-*0.4 and 5.6, molecular weight between 160 and 480, molar refractivity between 40 and 130 and a total number of atoms between 20 and 70. Lipinski’s rules [27] limit various parameters as molecular weight ≤ 500, H-bond acceptors ≤ 10, H-bond donors ≤ 5, calculated Log P (CLogP) ≤ 5. Ve-ber’s rules [6] limit only two criteria for oral bioavailability for the drug candidates and they are rotatable bonds ≤ 10 and polar surface area ≤ 140 A^02^ (or H-bond donors and acceptors ≤ 12). Egan’s rule [28] is defined by total polar surface area (TPSA) > 131.6Å or log P > 5.88. Muegge’s rule [44] defines two pharmacophore filter (PF1 and PF2) in addition to some additional rules which discriminate between drug and nondrug molecules. For finding the accurate compounds as leads we applied all of these rules for screening and it was found that only 7 compounds followed these rules. Because for making discrimination between non-drugs and drugs molecule, the rule of five cannot be employed. Physico-chemical parameters are may be required but are not effective features of a drug-like compound and Physico-chemical parameters cannot singly ascertain a drug molecule. For this, in the later phase of the drug discovery practice, other cogent aptitudes in crucial parameters like the half-life of analogs of lead molecules and oral bioavailability are accepted as standard among scientists. Thus, endeavors are given on other experimental data including membrane affinity, plasma exposure, and protein binding or aqueous solubility for correlating with counting parameters, for example, rule-of-five [29]. Based on the higher value of Pa compared to Pi, 6 compounds were selected which have better antimycobacterial activity. From DataWarrior software, it was found that only 5 compounds didn’t have any type of toxicity and had a good drug-likeness score. The term ‘drug-like’ is used by a little of variety by a number of authors [45–47]. Drug-likeness is defined commonly as a descriptor in a statistical way which is resultant from the catalogs of other compounds [29]. These 5 compounds are considered as leads. Leads categorized into three classes, first, low-affinity (>0.1 *µ*m) compounds with low ClogP and MW. By the optimization of pharmacokinetic profile and potency, these are turned into drugs. The second type of leads have high molecular weight and affinity and a number of drugs are included in this class. Finally, the last class of leads has drug-like ClogP (3-5) and MW (300-500) with low affinity [48].

### 3.3. Molecular Docking

Molecular docking paves the way for studying the mechanism of interaction between protein and compounds [49]. Higher efficacy between drug and protein could be indicated by the higher binding affinity [50]. After optimization by UCSF Chimera, leads and Benzodiazepine were docked with protein by using AutoDock Vina ver. 1.1.2.

Analysis of the docking result was done focusing mainly on three main points, i.e. binding energy (kcal/mol), number of H-bonds and interacting amino acids. In the case of molecular docking, the best binding energy between protein and ligand is indicated by the highest negative value and vice versa [51]. From Table 5, it is clear that compound with 13-Methoxymatrine (CHEMBL1672134) showed the highest binding affinity towards the target protein being analyzed. The interaction between protein and leads are shown in (two-dimensional) Figures 10, 11, 12, 13, 14 and 15 and (three-dimensional) Figures 16, 17, 18, 19, 20 and 21 and the interaction between protein and top lead is shown in Figures 22, 23 and 24. Interestingly, it was found that 13-Methoxymatrine had a good antimycobacterial as well as antibacterial activity. Table 6 shows the physical properties of the 13-Methoxymatrine compound. It is suggested that a ligand to be an ideal one should have the minimum docking score and must satisfy drug-likeness parameters [52]. Our selected ligand satisfied both of these criteria in accordance with having no mutagenic, tumorigenic, reproductive and irritant effect which follows Ghose’s rule, Lipinski’s rule, Veber’s rule and Egan’s rule containing a good antimycobacterial as well as antibacterial activity. So, 13-Methoxymatrine can be regarded as the final lead for becoming a potential drug for treating neuropathic pain by VGF.

**Table 5.**
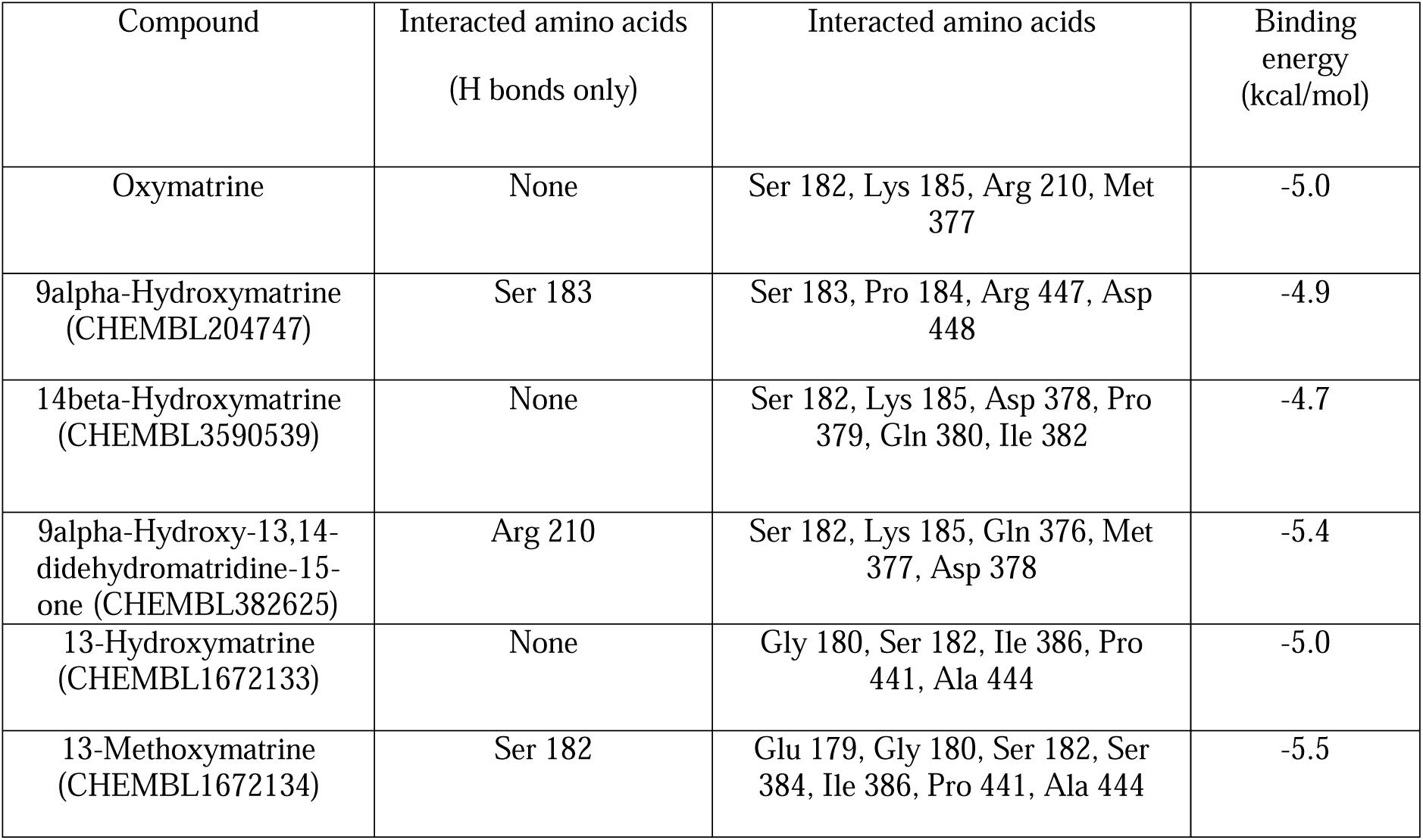
Binding energy and interacted amino acids of different compounds with the target protein.

| Compound | Interacted amino acids<br>(H bonds only) | Interacted amino acids | Binding energy<br>(kcal/mol) |
| --- | --- | --- | --- |
| Oxymatrine | None | Ser 182, Lys 185, Arg 210, Met 377 | -5.0 |
| 9alpha-Hydroxymatrine<br>(ChEMBL204747) | Ser 183 | Ser 183, Pro 184, Arg 447, Asp 448 | -4.9 |
| 14beta-Hydroxymatrine<br>(ChEMBL3590539) | None | Ser 182, Lys 185, Asp 378, Pro 379, Gln 380, Ile 382 | -4.7 |
| 9alpha-Hydroxy-13,14-didehydromatridine-15-one<br>(ChEMBL382625) | Arg 210 | Ser 182, Lys 185, Gln 376, Met 377, Asp 378 | -5.4 |
| 13-Hydroxymatrine<br>(ChEMBL1672133) | None | Gly 180, Ser 182, Ile 386, Pro 441, Ala 444 | -5.0 |
| 13-Methoxymatrine<br>(ChEMBL1672134) | Ser 182 | Glu 179, Gly 180, Ser 182, Ser 384, Ile 386, Pro 441, Ala 444 | -5.5 |

**Table 6.**
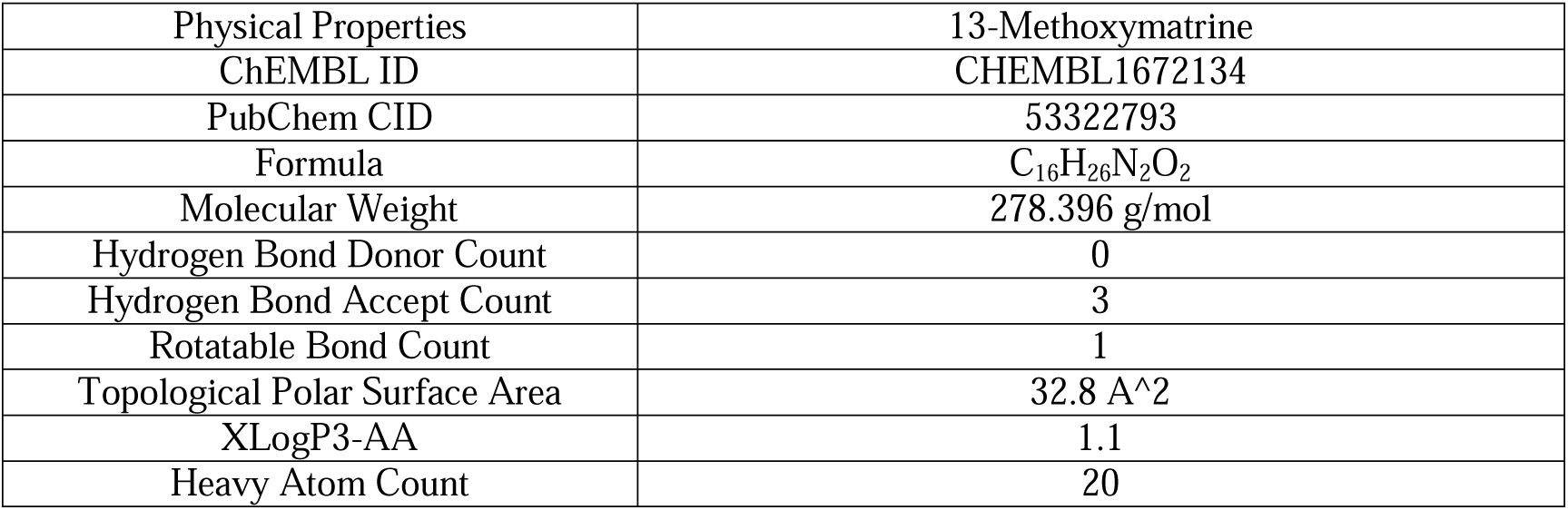
Physical properties of the top ranked molecule (13-Methoxymatrine).

**Fig. 10.**
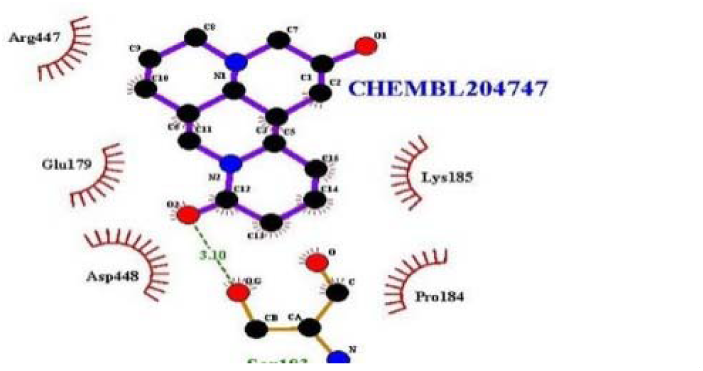
2D interaction between protein and 9alpha-Hydroxymatrine (CHEMBL204747).

**Fig. 11.**
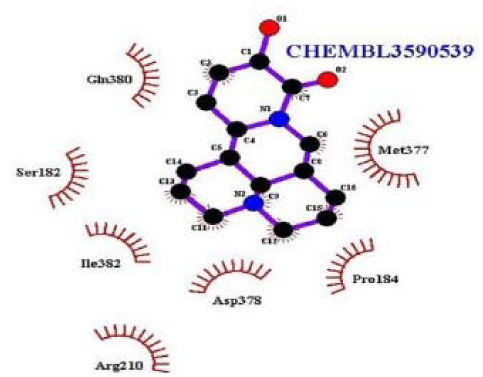
2D interaction between protein and 14beta-Hydroxymatrine (CHEMBL3590539).

**Fig. 12.**
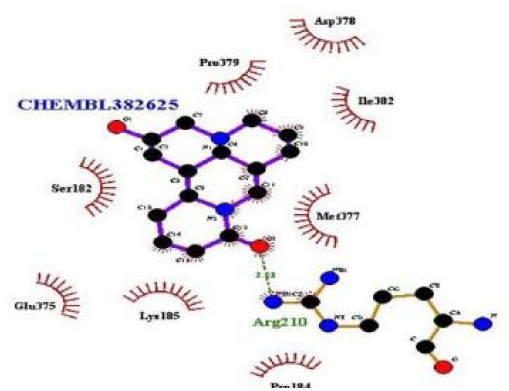
2D interaction between protein and 9alpha-Hydroxy-13,14-didehydromatridine-15-one (CHEMBL382625).

**Fig. 13.**
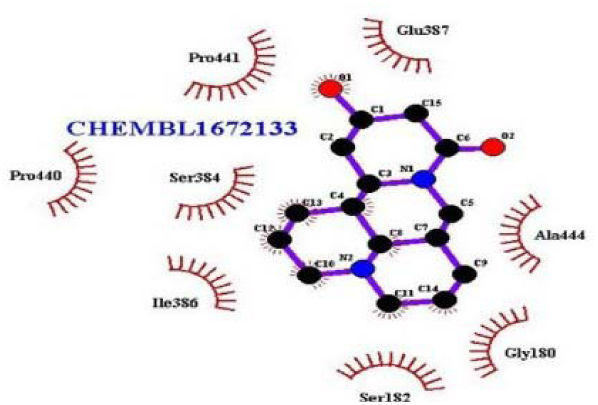
2D interaction between protein and 13-Hydroxymatrine (CHEMBL1672133).

**Fig. 14.**
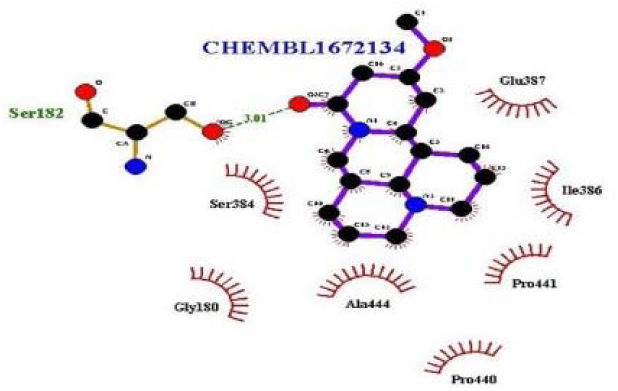
2D interaction between protein and 13-Methoxymatrine (CHEMBL1672134).

**Fig. 15.**
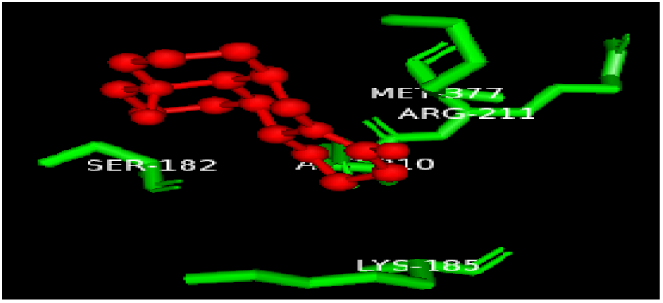
3D interaction between protein and Oxymatrine.

**Fig. 16.**
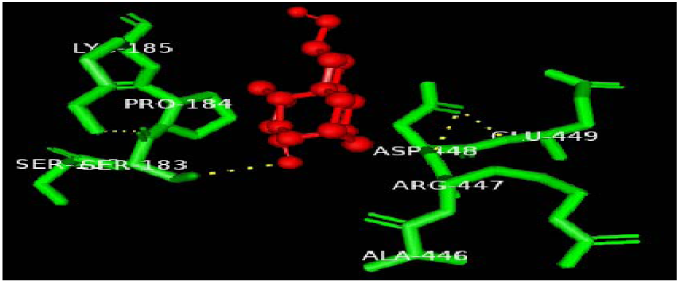
3D interaction between protein and 9alpha-Hydroxymatrine (CHEMBL204747).

**Fig. 17.**
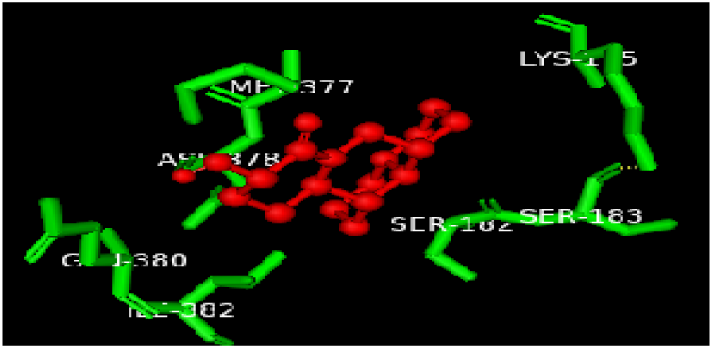
3D interaction between protein and 14beta-Hydroxymatrine (CHEMBL3590539).

**Fig. 18.**
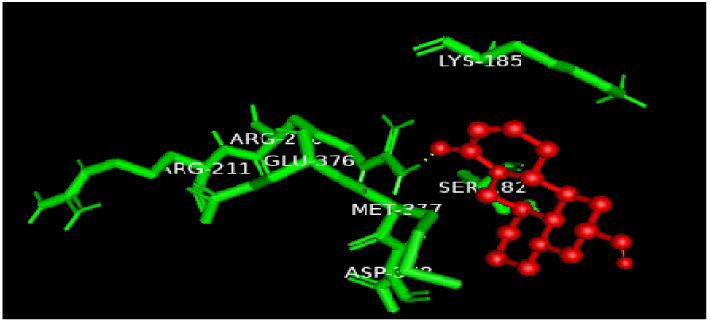
3D interaction between protein and 9alpha-Hydroxy-13,14-didehydromatridine-15-one (CHEMBL382625).

**Fig. 19.**
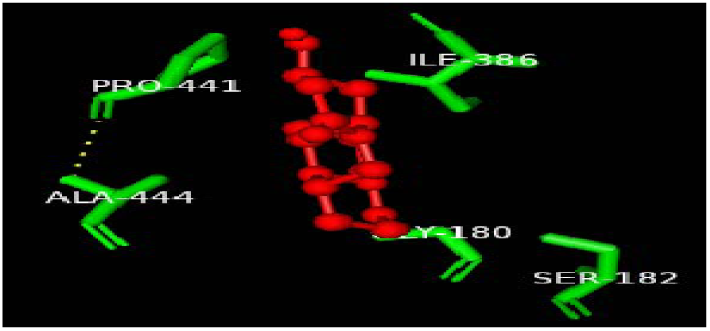
3D interaction between protein and 13-Hydroxymatrine (CHEMBL1672133).

**Fig. 20.**
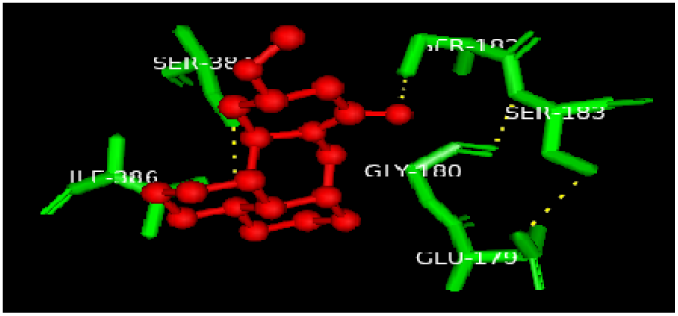
3D interaction between protein and 13-Methoxymatrine (CHEMBL1672134).

**Fig. 21.**
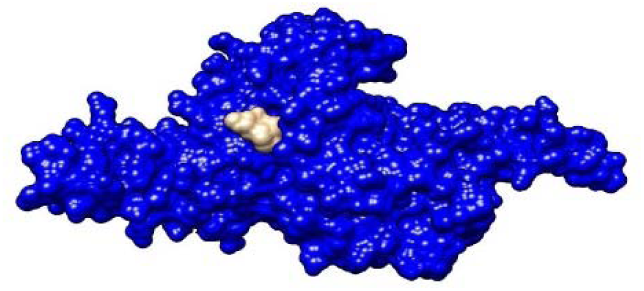
Interaction between protein (blue color) and top lead (13-Methoxymatrine) (grey color).

**Fig. 22.**
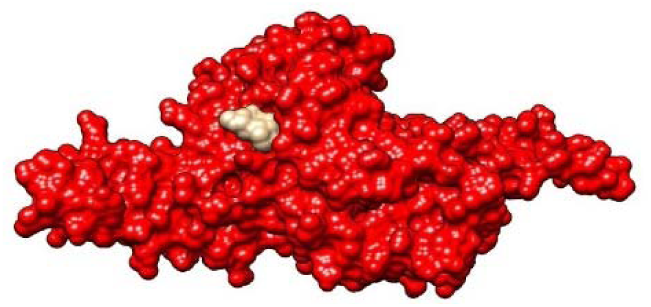
Interaction between protein (red color) and top lead (13-Methoxymatrine) (grey color).

**Fig. 23.**
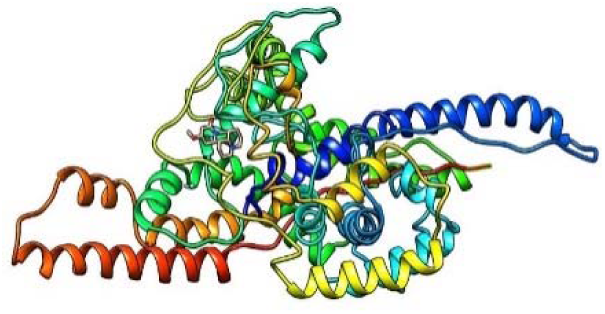
Interaction between protein (ribbon style) and top lead (13-Methoxymatrine) (wire-frame style).

### 3.4. Molecular properties and Oral toxicity prediction

To get the prediction of pharmacokinetics properties of the lead compound 13-Methoxymatrine and mother ligand Oxymatrine, the SwissADME server was used and results are shown in Table 7. The toxicity and ADME profile of the lead compound 13-Methoxymatrine and Oxymatrine were predicted by PreADMET server which is presented in Tables 8 and 9.

**Table 7.**
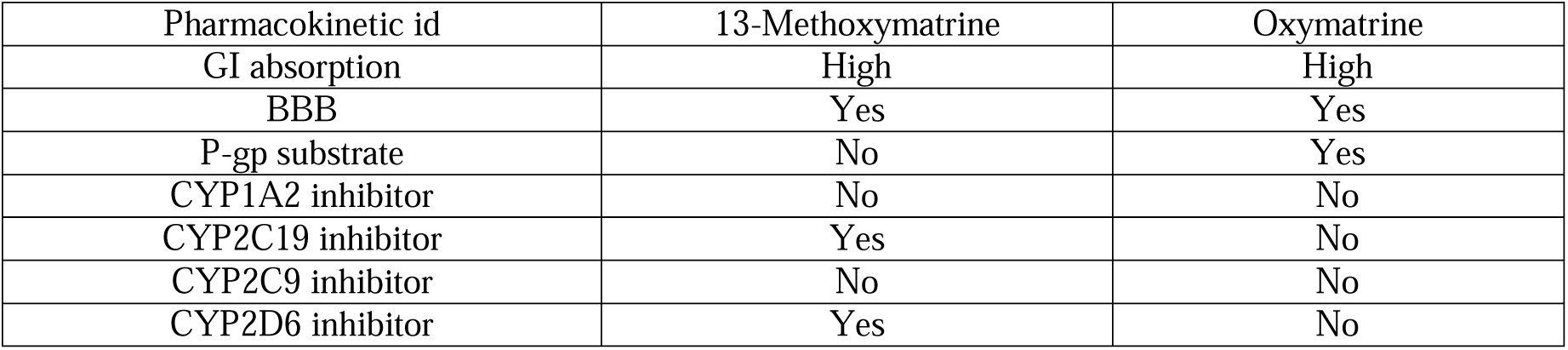
Predicted Pharmacokinetics profile by SwissADME server of lead compound (13-Methoxymatrine) and mother compound (Oxymatrine).

**Table 8.**
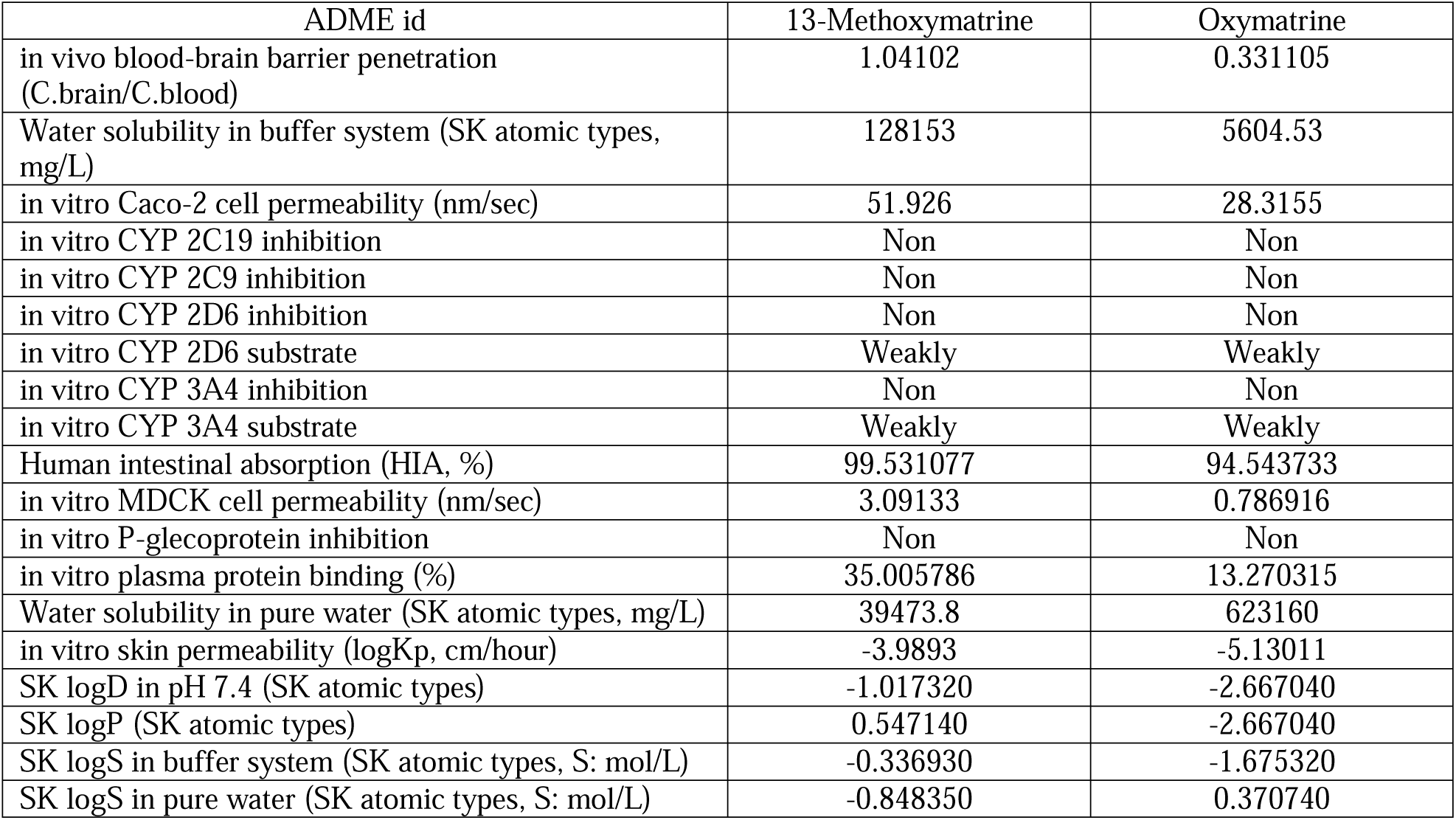
Predicted ADME profile by PreADMET server of lead compound (13-Methoxymatrine) and mother compound (Oxymatrine).

**Table 9.**
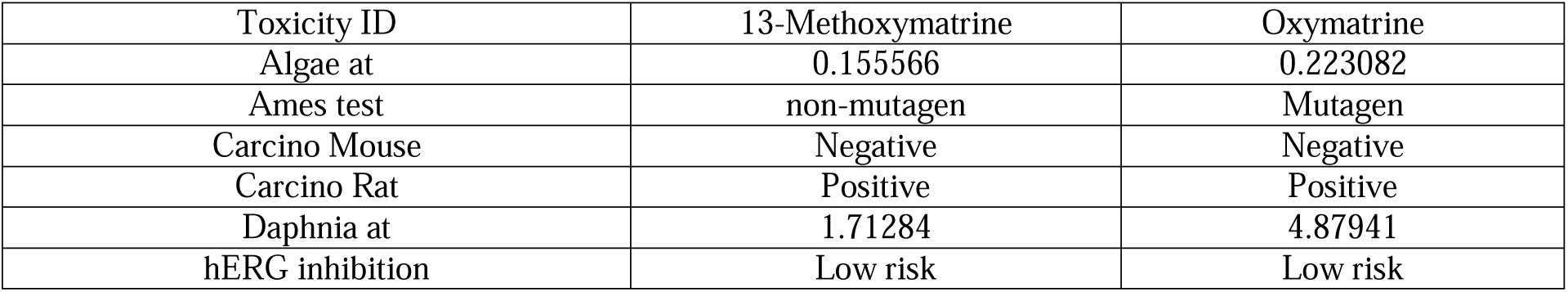
Predicted toxicity profile by PreADMET server of lead compound (13-Methoxymatrine) and mother compound (Oxymatrine).

To develop an effective drug BBB is a major obstacle and BBB inhibits entrance most of the drug in the brain [53]. It was found that about 98% of the drugs fail to reach the market only for their insufficient BBB permeability [54]. Our lead (13-Methoxymatrine) and mother compound (Oxymatrine) both have the ability of BBB penetration Tables 7 and 8. A drug should have the capability of passing through the biological membranes, most importantly through human intestinal mucosa, which has a crucial role in the drug development methodology and for getting regulatory approval [55] and for the approval of novel and effective oral drug, understanding the transportation system of the drug through the membrane of intestine (large and small) is also prerequisite [56–61] and our lead as well as mother drug also positive in this criteria. Another major barrier towards successful absorption of any drug candidate is whether it is able to act as a substrate for P-glycoprotein (P-gp) or not which is an efflux membrane transporter and limits cellular uptake of a drug [62]. Our proposed lead (13-Methoxymatrine) predicted as Non-P-gp substrate which demands some optimization while mother ligand Oxymatrine predicted as P-gp substrate. Due to the difficulty and costly nature of human intestinal permeability [63], Caco-2 or human colon carcinoma cell line is frequently used for screening permeability of drug candidates in the intestine [64]. But there is a considerable amount of conflicts between the Caco-2 permeability and measured the amount of HIA (human intestinal absorption) or permeability in the human intestine [65, 66]. In accordance with this, it is opined that biases or variability is introduced by in silico datasets as measurements of permeability is done from diverse laboratories and ambiguous information may be generated in terms of drug design [64]. It was predicted that our proposed lead (13-Methoxymatrine) has high Caco-2 permeability while mother compound (Oxymatrine) has medium Caco-2 permeability and for this, a wet lab experiment is deserved to quantify whether this lead has a good penetrability through Caco-2 or not.

In terms of metabolism of drugs, cytochrome P-450 has significant clinical drug-drug interactions. There are about 30 isoenzymes of cytochrome P-450 in human are identified and among them CYP1A2, CYP2C, CYP2D6, and CYP3A4 sub-family have a major role in the metabolism of drug [67].

Overall, our proposed lead (13-Methoxymatrine) and mother compound (Oxymatrine) are good enough in this case but demands some optimization as well as some wet lab experiments for ensuring the predicted value is accurate.

In terms of toxicity profile, our proposed lead is AMES non-mutagen. Our proposed lead (13-Methoxymatrine) predicted as non-mutagen crosses this barrier and mother ligand predicted as mutagen. Though, there are some critiques regarding AMES test. As in AMES test, rat liver S9 fraction is used and there are considerable amount of differences between rat and human metabolism regarding the mutagenic effect of chemicals and it is strongly recommended to use human liver S9 fraction instead of rat S9 in AMES test for assessing genotoxicity to humans [68]. For this, our lead (13-Methoxymatrine) also requires a wet lab experiment for evaluating whether it is toxic or non-toxic to human.

Failure of drug candidates during developmental stage accounts for about 20% due to toxic properties [69]. It is estimated that during the preclinical development stage, 54% failures occurred due to safety and toxicity concern which was responsible for US$1.8 billion/marketed drug [70]. It was found that the use of animals in trial phase were 100-200 million, 60-85 million and 50-100 million in 1970, 1993 and 2005 respectively [71]. A statistic showed that over 29 million animals were used for clinical trials only in North American and European countries [72]. In this situation, in silico methods for predicting the toxicity of a drug candidate can be used which will not only reduce time and cost but also save the lives of millions of animals [73]. PreADMET server predicted that our top lead (13-Methoxymatrine) and mother ligand (Oxymatrine) were negative in terms of carcinogenic in mouse as well as had low risk in terms of hERG inhibition.

## 4. CONCLUSION

In this current study, we have keyed out a potential drug candidate against VGF which is responsible for the regulation of metabolism, reproduction, gastric contractility, mood regulation, and peripheral neuropathic pain in human.

The compound 13-Methoxymatrine is an antagonist for VGF. This molecule has shown some good properties than the current mother ligand oxymatrine which is also a potential VGF inhibitor. The new molecule has also promising docking scores in the active site of the target VGF compared to oxymatrine. Additionally, this compound showed a good toxicity profile with some confusions in ADMET properties which require some wet lab experiments to verify the real scenario.

These analyses demonstrate that our proposed drug candidate can be a good choice of option for treating the peripheral neuropathic pain caused by VGF. Furthermore, evaluation in the wet lab is obligatory for getting regulatory approval of a drug candidate.

However, these findings provide a consolidated suggestion that our proposed lead can be a drug candidate and could be used as drug template for the treatment of peripheral neuropathic pain caused by VGF in human.

## Supporting information

List of tables

## 5. LIST OF ABBREVIATIONS

ADMET: absorption, distribution, metabolism, and excretion - toxicity in pharmacokinetics
AMES test: a test to determine the mutagenic activity of chemicals. BBB-blood brain barrier
BLAST: Basic Local Alignment Search Tool
CADD: Computer-aided drug discovery
ChEMBL: Biological database
COMT: Catechol-O-Methyltransferase
CYP2D6: Cytochrome P450 2D6
EMBL: European Molecular Biology Laboratory
HEK cells: Human embryonic kidney cells
HGMD: Human Gene Mutation Database
kDa: kilodaltons
KOR: κ-opioid receptor
MAOA: Monoamine Oxidase A
MDCK: Madin-Darby Canine Kidney (MDCK) cells
MOR: µ-opioid receptor
NCBI: National Centre for Biotechnology Information
OMTR: Oxymatrine
PDB: Protein Data Bank
PDBQT: Protein Data Bank, Partial Charge (Q), & Atom Type (T) format
PreADMET: web-based application for predicting ADME data
UCSC: University of California, Santa Cruz OPRM1,
UCSF: University of California, San Francisco

## 7. ACKNOWLEDGEMENTS

The authors are highly acknowledged and express their gratitude to late Dr. Md. Shamim Akhter for his high support and motivation.

