## Supplementary material for "A Comprehensive *In Silico* Study for the Identification of Therapeutic Target Against Peripheral Neuropathic Pain in humans": List of tables

1 Templates used for homology modeling of VGF	3
2 Ramachandran plot statistics for protein model	4
3 Root mean square deviation (RMSD) between VGF model and template (3j83.1. A).	5
4 Root mean square deviation (RMSD) between VGF model and template (2i1j)	5
5 Binding energy and interacted amino acids of different compounds with the target protein	7
6 Physical properties of the top ranked molecule (13-Methoxymatrine)	7
7 Predicted Pharmacokinetics profile by SwissADME server of lead compound (13-Methoxymatrine) and mother compound (Oxymatrine)…………………………………………………11
8 Predicted ADME profile by PreADMET server of lead compound (13-Methoxymatrine) and mother compound (Oxymatrine).	11
9 Predicted toxicity profile by PreADMET server of lead compound (13-Methoxymatrine) and mother compound (Oxymatrine).	11
