## Supplementary material for "A Comprehensive *In Silico* Study for the Identification of Therapeutic Target Against Peripheral Neuropathic Pain in humans": Table 1

**Table 1** Templates used for homology modeling of VGF

| PDB ID | DESCRIPTION | IDENTITY (%) | QUERY COVER (%) |
| --- | --- | --- | --- |
| 3j83 1A | Heptameric EspB Rosetta model guided by EM density | 34.62 | 52.5 |
| 2I1J A | Moesin from Spodoptera frugiperda at 2.1 angstroms resolution | 26.47 | 42.3 |
