## Supplementary material for "A Comprehensive *In Silico* Study for the Identification of Therapeutic Target Against Peripheral Neuropathic Pain in humans": Table 2

**Table 2** Ramachandran plot statistics for protein model

| Ramachandran plot statistics | Residues | % |
| --- | --- | --- |
| Residues in most favoured regions [A,B,L] | 387 | 87 |
| Residues in additional allowed regions [a,b,l,p] | 41 | 9.2 |
| Residues in generously allowed regions [~a,~b,~l,~p] | 10 | 2.2 |
| Residues in disallowed regions | 7 | 1.6 |
| Number of non-glycine and non-proline residues | 445 | 100 |
| Number of end-residues (excl. Gly and Pro) | 0 |  |
| Number of glycine residues (shown as triangles) | 34 |  |
| Number of proline residues | 66 |  |
| Total number of residues | 545 |  |
