## Supplementary material for "A Comprehensive *In Silico* Study for the Identification of Therapeutic Target Against Peripheral Neuropathic Pain in humans": Table 3

**Table 3** Root mean square deviation (RMSD) between VGF model and template (3j83.1.A)

| Name | RMSD Value |
| --- | --- |
| VGF Protein model | 1.169 angstroms; (across all 256 pairs) |
| 3j83.1.A |  |
