## Supplementary material for "A Comprehensive *In Silico* Study for the Identification of Therapeutic Target Against Peripheral Neuropathic Pain in humans": Table 5

**Table 5** Binding energy and interacted amino acids of different compounds with the target protein

| Compounds | Interacted amino acids  (H bonds only) | Interacted amino acids | Binding energy (kcal/mol) |
| --- | --- | --- | --- |
| Oxymatrine or CHEMBL458337 | None | Ser 182, Lys 185, Arg 210, Met 377 | -5.0 |
| CHEMBL204747 | Ser 183 | Ser 183, Pro 184, Arg 447, Asp 448 | -4.9 |
| CHEMBL3590539 | None | Ser 182, Lys 185, Asp 378, Pro 379, Gln 380, Ile 382 | -4.7 |
| CHEMBL382625 | Arg 210 | Ser 182, Lys 185, Gln 376, Met 377, Asp 378 | -5.4 |
| CHEMBL1672133 | None | Gly 180, Ser 182, Ile 386, Pro 441, Ala 444 | -5.0 |
| CHEMBL1672134 | Ser 182 | Glu 179, Gly 180, Ser 182, Ser 384, Ile 386, Pro 441, Ala 444 | -5.5 |
