## Supplementary material for "A Comprehensive *In Silico* Study for the Identification of Therapeutic Target Against Peripheral Neuropathic Pain in humans": Table 6

**Table 6** Physical properties of the top ranked molecule (13-Methoxymatrine)

| Physical Properties | 13-Methoxymatrine |
| --- | --- |
| ChEMBL ID | CHEMBL1672134 |
| PubChem CID | 53322793 |
| Formula | C_16_H_26_N_2_O_2_ |
| Molecular Weight | 278.396 g/mol |
| Hydrogen Bond Donor Count | 0 |
| Hydrogen Bond Accept Count | 3 |
| Rotatable Bond Count | 1 |
| Topological Polar Surface Area | 32.8 A^2 |
| XLogP3-AA | 1.1 |
| Heavy Atom Count | 20 |
