## Supplementary material for "A Comprehensive *In Silico* Study for the Identification of Therapeutic Target Against Peripheral Neuropathic Pain in humans": Table 7

**Table 7** Predicted Pharmacokinetics profile by SwissADME server of lead compound (13-Methoxymatrine) and mother compound (Oxymatrine)

| Pharmacokinetic id | CHEMBL1672134 | Oxymatrine |
| --- | --- | --- |
| GI absorption | High | High |
| BBB | Yes | Yes |
| P-gp substrate | No | Yes |
| CYP1A2 inhibitor | No | No |
| CYP2C19 inhibitor | Yes | No |
| CYP2C9 inhibitor | No | No |
| CYP2D6 inhibitor | Yes | No |
