## Supplementary material for "A Comprehensive *In Silico* Study for the Identification of Therapeutic Target Against Peripheral Neuropathic Pain in humans": Table 8

**Table 8** Predicted ADME profile by PreADMET server of lead compound (13-Methoxymatrine) and mother compound (Oxymatrine).

| ADME id | CHEMBL1672134 | Oxymatrine |
| --- | --- | --- |
| in vivo blood-brain barrier penetration (C.brain/C.blood) | 1.04102 | 0.331105 |
| Water solubility in buffer system (SK atomic types, mg/L) | 128153 | 5604.53 |
| in vitro Caco-2 cell permeability (nm/sec) | 51.926 | 28.3155 |
| in vitro CYP 2C19 inhibition | Non | Non |
| in vitro CYP 2C9 inhibition | Non | Non |
| in vitro CYP 2D6 inhibition | Non | Non |
| in vitro CYP 2D6 substrate | Weakly | Weakly |
| in vitro CYP 3A4 inhibition | Non | Non |
| in vitro CYP 3A4 substrate | Weakly | Weakly |
| Human intestinal absorption (HIA, %) | 99.531077 | 94.543733 |
| in vitro MDCK cell permeability (nm/sec) | 3.09133 | 0.786916 |
| in vitro P-glecoprotein inhibition | Non | Non |
| in vitro plasma protein binding (%) | 35.005786 | 13.270315 |
| Water solubility in pure water (SK atomic types, mg/L) | 39473.8 | 623160 |
| in vitro skin permeability (logKp, cm/hour) | -3.9893 | -5.13011 |
| SK logD in pH 7.4 (SK atomic types) | -1.017320 | -2.667040 |
| SK logP (SK atomic types) | 0.547140 | -2.667040 |
| SK logS in buffer system (SK atomic types, S: mol/L) | -0.336930 | -1.675320 |
| SK logS in pure water (SK atomic types, S: mol/L) | -0.848350 | 0.370740 |
