## Supplementary material for "A Comprehensive *In Silico* Study for the Identification of Therapeutic Target Against Peripheral Neuropathic Pain in humans": Table 9

**Table 9** Predicted toxicity profile by PreADMET server of lead compound (13-Methoxymatrine) and mother compound (Oxymatrine).

| Toxicity ID | CHEMBL1672134 | Oxymatrine |
| --- | --- | --- |
| Algae at | 0.155566 | 0.223082 |
| Ames test | non-mutagen | Mutagen |
| Carcino Mouse | Negative | Negative |
| Carcino Rat | Positive | Positive |
| Daphnia at | 1.71284 | 4.87941 |
| hERG inhibition | Low risk | Low risk |
